# *H19* is a PERK-regulated long non-coding RNA that fine-tunes UPR signalling and inhibits endoplasmic reticulum stress-induced cell death

**DOI:** 10.1101/2025.10.17.683008

**Authors:** Wen Liu, Ananya Gupta, Michael Kerin, Sanjeev Gupta

## Abstract

The endoplasmic reticulum (ER) responds to stimuli that disrupt its homeostasis by activating a signalling network known as unfolded protein response (UPR), that restores cellular balance and determines cell fate through three key sensors: IRE1, PERK, and ATF6. Emerging evidence suggests that the UPR regulates the expression of numerous long non-coding RNAs (lncRNAs), which play critical roles in modulating stress responses. Here we show that expression of lncRNA *H19* is downregulated in response to ER stress in breast cancer cells. Using genetic and pharmacological approaches, we demonstrate that this downregulation is primarily mediated by the PERK arm of the UPR. Specifically, knockdown or chemical inhibition of PERK compromised the ER stress-mediated *H19* repression, while PERK activation significantly reduced *H19* expression. *H19* overexpression promotes the optimal activation of ATF6 and PERK pathways, while it attenuates the signalling by IRE1-XBP1 axis of the UPR. Bioinformatic analyses across multiple breast cancer cohorts revealed that high *H19* expression correlates with poor prognosis, particularly in basal-like subtypes. Furthermore, in triple-negative breast cancer (TNBC) cells, *H19* was found to provide resistance to ER stress-induced apoptosis. Collectively, our findings establish *H19* as a key regulatory node in the UPR network, where PERK-mediated repression of *H19* shapes cellular outcomes under ER stress. These insights position *H19* as a potential therapeutic target in breast cancer, especially in aggressive subtypes such as TNBC.

## Introduction

Unfolded protein response (UPR) is an evolutionarily conserved pathway in eukaryotic cells comprising three main sensors - inositol requiring enzyme1α (IRE1α), activating transcription factor-6 (ATF6), and protein kinase RNA-like endoplasmic reticulum kinase (PERK). Under normal conditions the activity of UPR sensors is kept under control by interaction with molecular chaperone glucose-regulated protein 78 (GRP78, also known as (BiP /HSPA5) in the lumen of endoplasmic reticulum (ER). Upon mild ER stress, a condition in which unfolded or misfolded proteins accumulate in the ER lumen, GRP78 dissociates from PERK enabling its activation by dimerisation and autophosphorylation. Activated PERK phosphorylates eIF2α to reduce global protein synthesis, alleviating the ER’s protein-folding burden[1]. Concurrently, PERK activation promotes the selective translation of activating transcription factor 4 (ATF4), a transcription factor that upregulates genes involved in amino acid metabolism, redox homeostasis, and ER chaperone expression. Similarly, dissociation of GRP78 from IRE1 leads to its activation by dimerisation and auto-phosphorylation leading to activating of its endoribonuclease domain, which catalyses the unconventional splicing of X-box binding protein-1 (XBP1) mRNA to produce the potent transcription factor spliced XBP1 (XBP1s). Consequently, XBP1s moves to the nucleus and drives the expression of genes involved in protein folding, ER-associated degradation (ERAD), and lipid biosynthesis[2]. In parallel, upon dissociation of GRP78 the full-length ATF6 (ATF6p90) translocates from the ER to the Golgi apparatus, where it undergoes proteolytic cleavage to release its active cytoplasmic domain, ATF6p50. The cytosolic ATF6p50 functions as a transcription factor, upregulating ER chaperones such as GRP78 and components of the ERAD pathway[3]. Together, these adaptive mechanisms enhance the cell’s capacity to manage stress and restore ER homeostasis. However, under prolonged or severe ER stress, ATF4 upregulates genes involved in redox homeostasis, such as C/EBP homologous protein (CHOP). CHOP is a pro-apoptotic transcription factor that promotes apoptosis by downregulating anti-apoptotic proteins such as B-cell lymphoma 2 (Bcl-2) and upregulating pro-apoptotic proteins like Bcl-2-interacting mediator of cell death (BIM) and death receptor 5 (DR5) [4, 5]. Additionally, sustained PERK activation can lead to the accumulation of reactive oxygen species, further exacerbating cellular damage and driving cell death[6]. IRE1 can switch from XBP1 splicing to regulated IRE1-dependent decay, reducing the synthesis of proteins entering the ER, but excessive mRNA decay can lead to the loss of critical proteins, ultimately promoting cell death[7]. Overall, UPR plays a dual role in normal cells: it promotes cell survival by enhancing protein folding and reducing ER stress under mild conditions, but it switches to inducing apoptosis under severe or prolonged stress, thereby eliminating damaged cells and maintaining tissue homeostasis.

Long non-coding RNAs (lncRNAs) are a class of non-coding RNAs longer than 200 nucleotides that do not encode proteins but play key regulatory roles in gene expression. LncRNAs exert their regulatory effects through interactions with DNA, RNA, and proteins: in the nucleus, they can modify chromatin or influence transcription by recruiting transcription factors or chromatin remodelling enzymes; in the cytoplasm, they may act as competing endogenous RNAs (ceRNAs), sequester regulatory molecules, or modulate signalling pathways through protein binding. Emerging evidence links ncRNAs to UPR signalling, and during the past few years, work from several groups has revealed that all three branches of the UPR regulate specific subsets of lncRNAs. For example, CHOP regulates the expression of *lnc-MGC* and lncRNA golgin A2 pseudogene 10 (*GOLGA2P10*); *MALAT1* expression is regulated by IRE1 and PERK pathways [8, 9]. The outcome of UPR-regulated ncRNA expression is the fine-tuning of the UPR signalling to modulate cellular adaptation to stress and regulation of cell fate.

The long non-coding RNA *H19* (lncRNA *H19*) is a highly conserved and multifunctional RNA molecule that plays significant roles in development, disease, and gene regulation. *H19* is a maternally expressed gene located on chromosome 11p15.5. While *H19* is highly expressed during embryogenesis and generally silenced in adult tissues, its reactivation has been observed in numerous cancers, where it plays diverse and often oncogenic roles[10, 11]. *H19* is upregulated in many solid tumours, including breast, liver, colorectal, gastric, bladder, and lung cancers[12–14] and contributes to multiple hallmarks of cancer. Conversely, under specific conditions or in certain tumour types, *H19* has also been suggested to exert tumour-suppressive roles, reflecting a complex and context-specific regulatory landscape[15, 16]. Mechanistically, *H19* functions through several pathways: acting as a ceRNA, sponging tumour-suppressive miRNAs (e.g., *miR-200*, *miR-138*, and *miR-29a*)[17–22]; serving as a precursor for miR-675, which modulates targets involved in tumour progression[12, 23–25]; modulating gene expression through interactions with epigenetic regulators (e.g., *EZH2*, *MBD1*)[26–29]. Beyond its role in tumour progression, *H19* is increasingly implicated in cancer therapy resistance[30, 31]. Overexpression of *H19* has been associated with reduced sensitivity to chemotherapy agents (cisplatin, doxorubicin), and radiotherapy, often by inhibiting apoptosis, promoting autophagy, or enhancing stress adaptation[32–36]. *H19* can regulate drug resistance genes or affect pathways like phosphoinositide 3-kinase/protein kinase B (PI3K/AKT), epithelial–mesenchymal transition (EMT), NF- κB, and mTOR, which are crucial for survival under therapeutic stress[37–40]. Given that both *H19* and the UPR are crucial for cellular adaptation and survival under stress, their potential interaction may represent a key axis in tumour resilience and therapy resistance. Despite wealth of knowledge about function of *H19* in human cancers not much is known about mechanisms regulating its expression.

In this study, we evaluated the role of *H19* during conditions of UPR. During conditions of UPR, the expression of both *H19* and *miR-675-5p* (miRNA generated from *H19*) was decreased in a PERK-dependent manner. The ectopic expression of *H19* modulated the optimal activation of ER stress sensors during conditions of UPR. Higher expression of *H19* was associated with poor overall survival (OS) in basal-like breast cancer. Finally, we show that *H19* inhibits ER stress-induced cell death. In summary, our results suggest that downregulation of *H19* by stressful conditions in tumour microenvironment may contribute to cancer progression by regulating UPR signalling and cell fate.

## Results

### Downregulation of *H19* during conditions of ER stress

To understand the upstream regulators of *H19* expression, we analysed the gene expression profile [GSE63252: Affymetrix microarrays in two breast cancer cell lines (T47D and MDA-MB-231) under treated (hypoxia and glucose deprivation) or untreated conditions with control and XBP1 knockdown] and observed a marked downregulation of *H19* expression upon knockdown of XBP1. To experimentally determine whether UPR regulates the expression of *H19*, we treated breast cancer cell lines MCF7 and T47D with classical UPR inducers: brefeldin A (BFA) and thapsigargin (TG). BFA is inhibitor of anterograde transport from the ER to the Golgi apparatus and TG is an inhibitor of sarcoplasmic/endoplasmic reticulum calcium ATPase . Both agents disrupt protein homeostasis by promoting the accumulation of unfolded or misfolded proteins in the ER, thereby triggering ER stress and activating the UPR. After 24 hours of treatment with BFA or TG, expression of *GRP78*—a canonical UPR target gene - was significantly upregulated, confirming successful activation of UPR. In parallel, *H19* expression was significantly repressed in both MCF7 and T47D cells (Fig. 1A-B). A time-course analysis further revealed a progressive decrease in *H19* expression following BFA treatment in MCF7 cells (Fig. 1C). To determine whether this repression of *H19* was restricted to breast cancer cells, we examined *H19* expression in HEK293T cells under ER stress conditions. Consistent with our earlier observations, BFA treatment induced GRP78 expression and caused a time-dependent reduction in *H19* levels (Fig. 1D). Together, these results demonstrate that *H19* is consistently downregulated in response to ER stress across multiple cell types and in response to different UPR-inducing agents. LncRNAs can be processed to generate functional microRNAs. Indeed, *H19* serves as the precursor transcript for *miR- 675-5p* and *miR-675-3p*, both of which are processed from the first exon of the *H19* transcript (Supplementary Figure 1A). The *miR-675-5p* (guide strand) accumulates at a higher level than *miR-675-3p* (passenger strand) (Supplementary Figure 1B). To explore the co-regulation of *H19* and its processed miRNAs under ER stress, we quantified the expression levels of *miR-675-5p* and *miR-675-3p* using microRNA quantitative PCR (miRNA-qPCR) in the same RNA samples previously analysed for *H19* expression (Supplementary Figure 1C). In MCF7 cells, *miR-675-5p* expression was significantly reduced by both BFA and TG treatment (Supplementary Figure 1C). These results suggest that during conditions of UPR, the expression of *miR-675-5p* (miRNA guide strand) and its primary transcript *H19* are downregulated in a coordinated manner.

**Figure 1:**
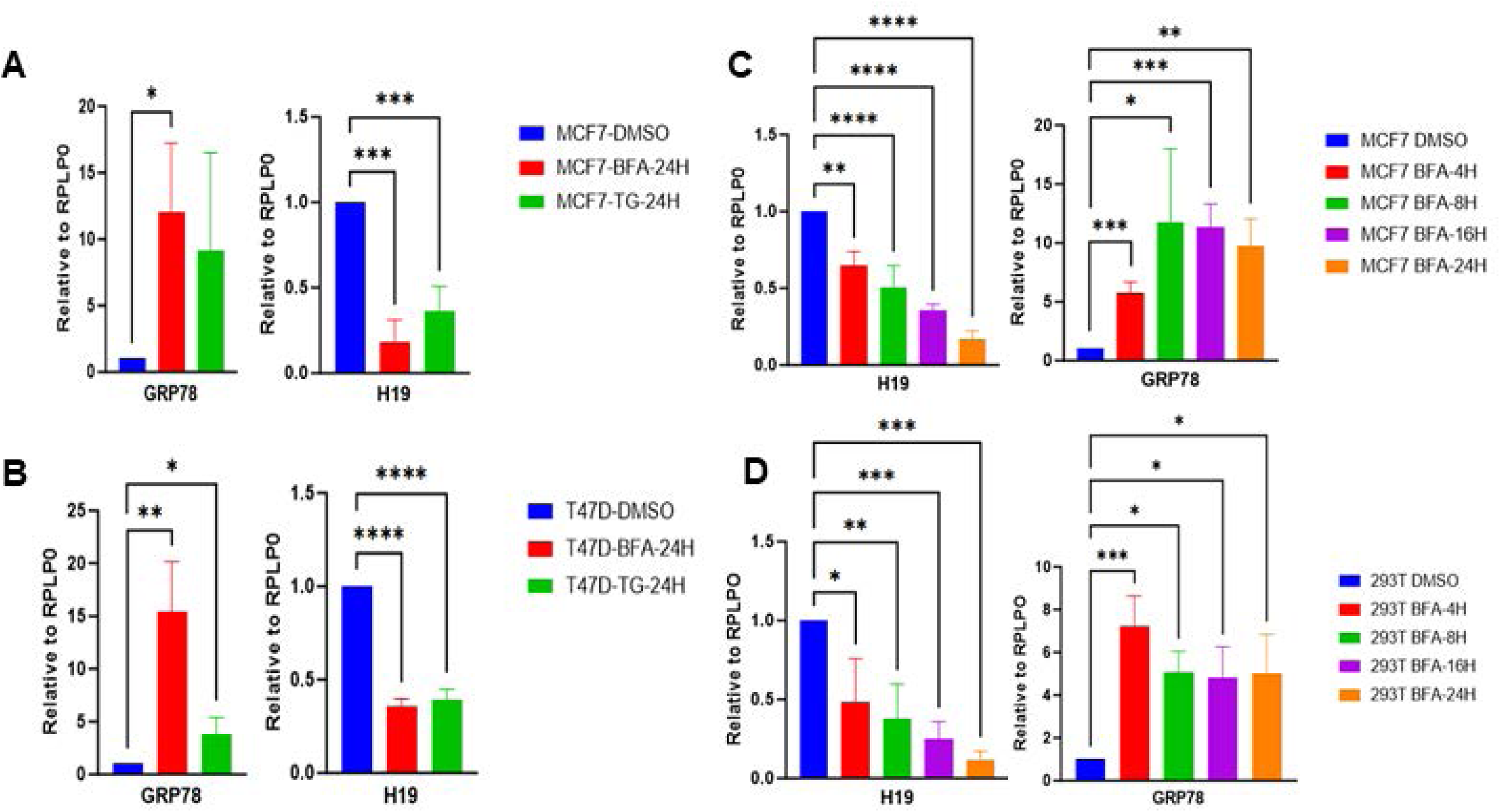
Expression of lncRNA *H19* is downregulated by UPR. (A–B) MCF7 (A) and T47D (B) cells were treated with either vehicle (DMSO) or thapsigargin (TG, 1 µM) and brefeldin A (BFA, 0.5 µg/ml) for 24 h. (C-D) Time-course analysis of BFA treatment in MCF7 (C) and HEK293T cells (D). Expression levels of *GRP78* and lncRNA *H19* were measured by RT-qPCR and normalized to *RPLP0*. Error bars represent the mean ± SD from three independent experiments performed in triplicates. Statistical significance was determined using an unpaired t-test compared with DMSO-treated cells. *P < 0.05, **P < 0.01, ***P < 0.001, ***P < 0.0001.

### Downregulation of *H19* expression is PERK-dependent

Next, we investigated the role of the UPR signalling pathways - PERK, IRE1, and ATF6 - in the downregulation of *H19* expression. To this end, we used MCF7 control (MCF7 PLKO) and UPR signalling-deficient MCF7 subclones (MCF7-XBP1 KD, MCF7-PERK KD and MCF7-ATF6 KD). The generation and characterisation of these subclones has been previously reported[41, 42]. The expression of PERK, XBP1, and ATF6 protein were significantly reduced (basal and BFA-treated) in respective knockdown subclones, confirming the knockdown of the target protein (Fig. 2A). Our results suggest a crosstalk between UPR arms, as ATF6 knockdown led to a reduction in XBP1 expression (Fig. 2A), which is consistent with previous reports showing that ATF6 regulates XBP1[43]. Notably, RT-qPCR analysis revealed that PERK knockdown partially compromised the down-regulation of *H19* during UPR (Fig. 2B).

**Figure 2:**
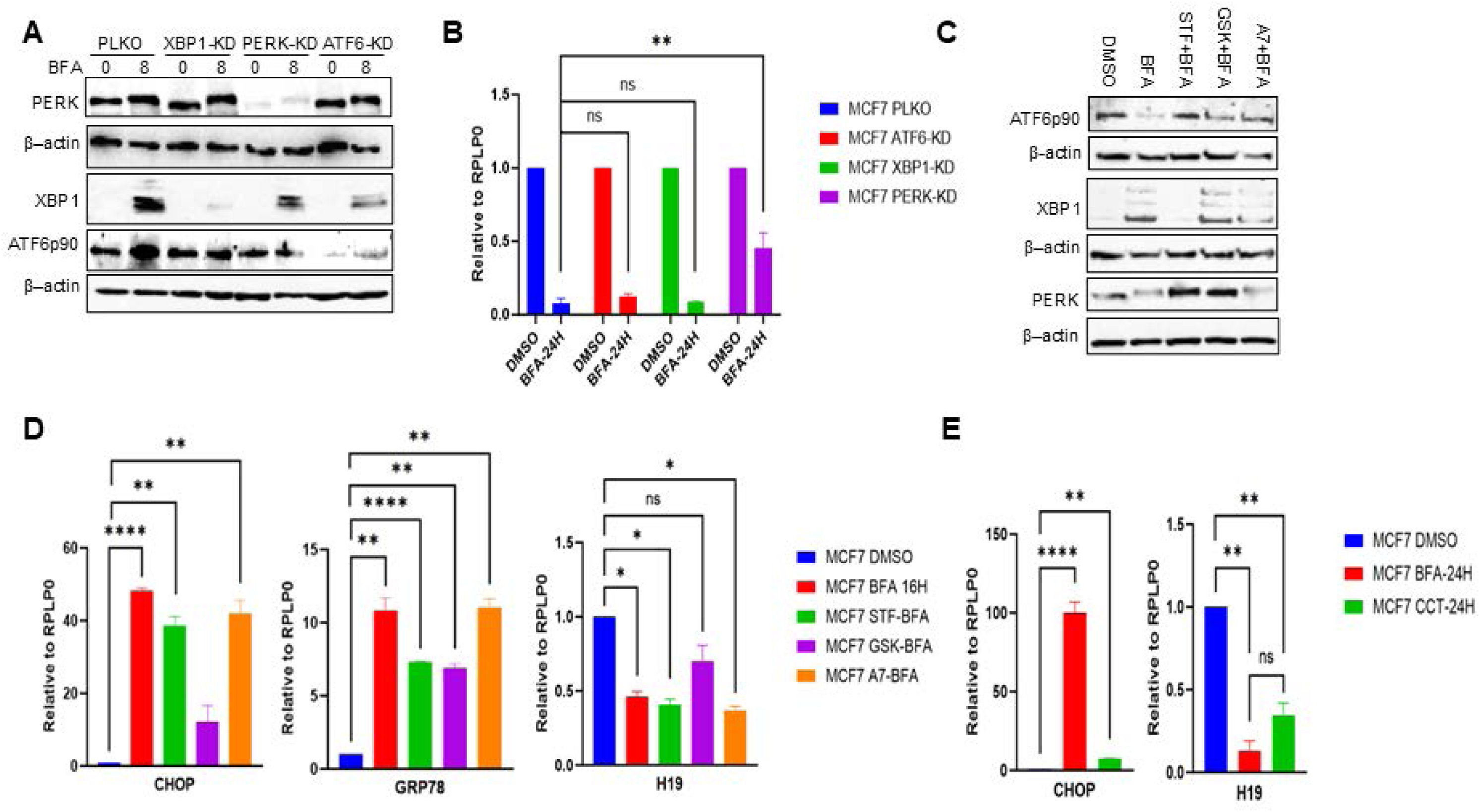
Downregulation of *H19* during ER stress is dependent on the PERK pathway. (A) MCF7 PLKO, MCF7 XBP1-KD, MCF7 PERK-KD, and MCF7 ATF6-KD subclones were treated with either vehicle (DMSO) or BFA (0.5 µg/ml) for 8 hours. Whole cell lysates were analysed by western blot using antibodies against ATF6, XBP1, PERK and β-Actin. (B) Cells were treated as in A, and expression levels of lncRNA *H19* were quantified by RT-qPCR normalising against *RPLP0*. (C) MCF7 cells were treated with BFA (0.5 µg/ml) in absence and presence of ATF6 inhibitor Ceapin A7 (A7, 10 µM), the IRE1α inhibitor STF083010 (STF, 50 µM), or the PERK inhibitor GSK2606414 (GSK, 5 µM). Whole cell lysates were analysed by western blot analysis using antibodies against ATF6, XBP1, PERK and β-Actin. (D) Cells were treated as in C, and total RNA was analysed by RT-qPCR for expression of *CHOP*, *GRP78*, and *H19* normalising against *RPLP0*. (E) MCF7 cells were treated with BFA (0.5 µg/ml) or the PERK activator CCT020312 (CCT, 5 µM) for 24 hours. The expression levels of *CHOP* and *H19* were quantified by RT-qPCR. Error bars represent the mean ± SD from three independent experiments performed in triplicate. Statistical significance was determined using an unpaired t-test compared with DMSO-treated cells. *P < 0.05, **P < 0.01, ***P < 0.001, ***P < 0.0001.

To further verify the role of PERK signalling in *H19* downregulation during UPR, we used chemical inhibitors targeting each UPR arm. MCF7 cells were treated with either a vehicle control (DMSO) or BFA in combination with ATF6 inhibitor (Ceapin A7), IRE1α inhibitor (STF083010), or PERK inhibitor (GSK2606414). Western blot analysis confirmed a reduction in the production of XBP1s with STF083010, decreased proteolytic processing of ATF6 with Ceapin A7 and decreased phosphorylation of PERK with GSK2606414 (Fig. 2C). Notably, inhibition of the ATF6 arm also led to a reduction in XBP1s protein levels, further supporting the existence of crosstalk between these pathways (Fig. 2C). Consistently, RT-qPCR analysis demonstrated that chemical inhibition of the PERK significantly compromised the downregulation of *H19* and upregulation of CHOP upon BFA treatment of MCF7 (Fig. 2D).

Next, we utilised CCT020312, a selective activator of the PERK pathway, to further investigate its role in *H19* repression. MCF7 cells were treated with CCT020312, and BFA was used as a positive control. First, we assessed the effect of CCT020312 on the induction of CHOP, a downstream target of PERK. As expected, CCT020312 treatment led to a significant upregulation of CHOP, confirming the specific activation of the PERK pathway (Fig. 2E). Following PERK activation, we observed a significant repression of *H19* in both CCT020312- and BFA-treated cells as compared to vehicle-treated controls. Moreover, the degree of *H19* repression in CCT020312-treated cells was not comparable to that observed in BFA-treated cells (Fig. 2E). Collectively, these findings suggest that *H19* repression during UPR is PERK-dependent.

### *H19* modulates activation of UPR signalling and cell fate during ER stress

Next, we determined the effect of *H19* on activation of three branches of the UPR. To investigate the effects of *H19* on UPR activation, we used HEK293T cells, which are well-suited for reporter assays due to their high transfection efficiency. HEK293T cells were transiently transfected with either a control vector or an *H19*-overexpressing plasmid. After 48 hours, RT-qPCR analysis showed a significant reduction in the Ct value for *H19* in HEK293T cells transfected with the *H19*-overexpressing plasmid as compared to control, indicating robust *H19* overexpression (Fig. 3A). In the same samples the Ct values for the housekeeping gene *RPLP0* were comparable, confirming normalization consistency (Fig. 3A). Next, we employed the *H19*-overexpressing plasmid and its control in subsequent experiments to assess the role of *H19* in UPR signalling.

**Figure 3:**
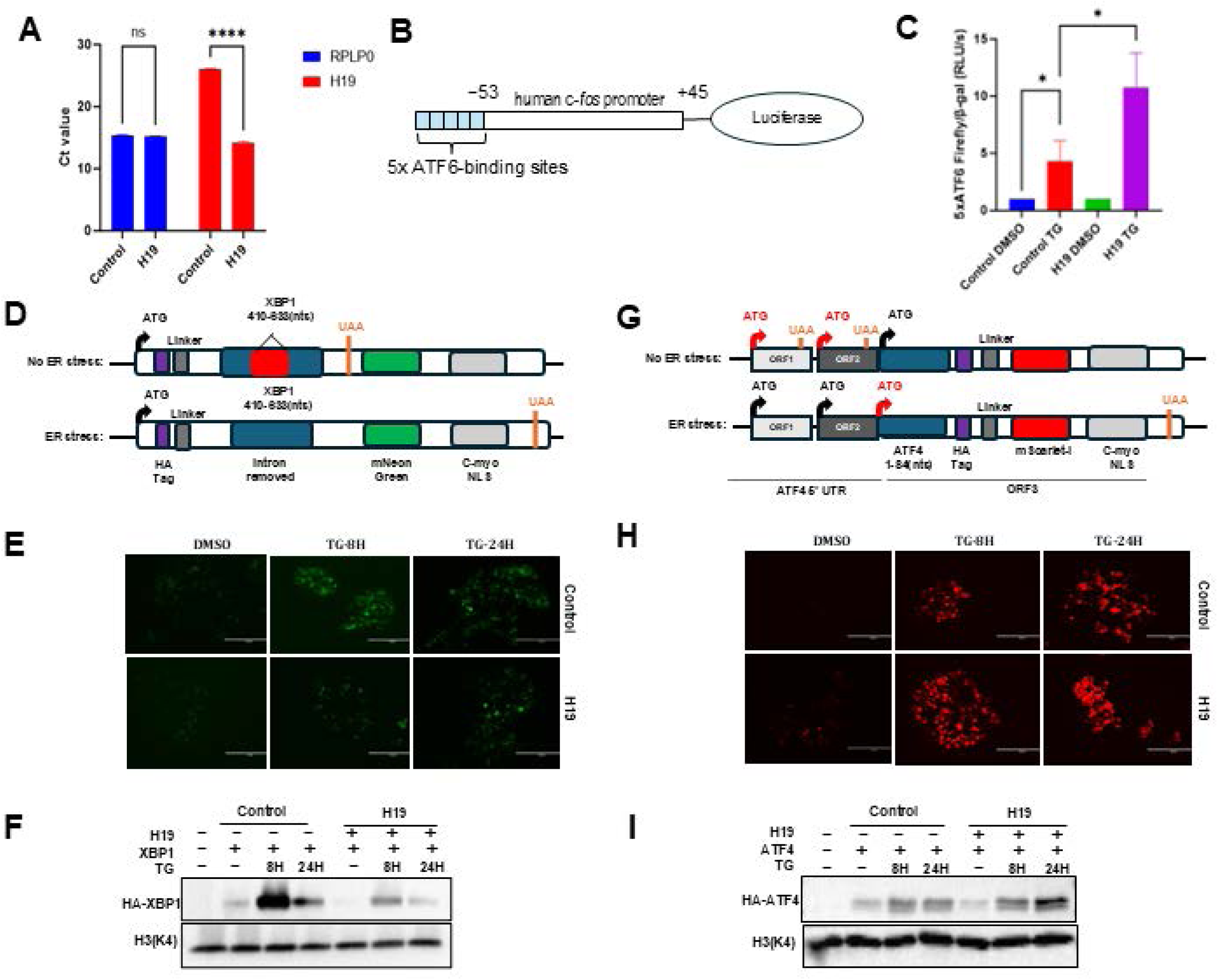
Effect of *H19* on activation of UPR signalling pathway. (A) RT-qPCR analysis of *H19* expression in HEK293T cells transfected with either H19-overexpressing plasmid (*H19*) or control vector (plenti4/V5) for 48 hours. (B) Schematic representation of the ATF6-luciferase reporter plasmid. (C) MCF7 cells were co-transfected with the *H19*-overexpressing plasmid or control vector, ATF6-luciferase reporter, and β-galactosidase plasmid. After 16 hours, cells were treated with thapsigargin (TG, 1 µM) for 24 hours. ATF6 transcriptional activity was assessed via firefly luciferase activity normalized to β-gal expression. (D–I) HEK293T cells were co-transfected with *H19*-overexpressing plasmid or control vector, and either the HA-tagged XBP1 reporter plasmid (D–F) or ATF4 reporter plasmid (G–I). (D, G) Diagrams of reporter plasmid structures. (E, H) GFP (XBP1) and RFP (ATF4) fluorescence signals recorded at 8 and 24 hours post-treatment with TG (1 µM). (F, I) Western blot analysis of HA-tagged XBP1 and ATF4 proteins, respectively, normalized to H3(K4).

To evaluate the impact of *H19* on the ATF6 signalling pathway, we used a synthetic luciferase reporter construct containing five tandem ATF6-binding elements (Fig. 3B) [44]. Luciferase reporter assays showed a significant increase in ATF6 transcriptional activity following TG treatment as compared to DMSO controls, confirming the responsiveness of the ATF6 reporter to ER stress. Importantly, *H19* overexpression further amplified TG-induced ATF6 reporter activity (Fig. 3C), indicating that *H19* enhances ATF6 signalling under ER stress conditions. To assess the role of *H19* in the IRE1-XBP1 signalling arm, we used a IRE1 activity reporter comprising of 410–633 nucleotide sequence of the *XBP1* cDNA, having a 26 bp intron that is spliced by RNase activity of activated IRE1. The splicing of the 26 bp intron leads to a change in reading frame that leads to translation of HA-tagged mNeonGreen protein fused with the c-myc NLS sequence, allowing the detection of IRE1 pathway activity by quantifying GFP fluorescence and/or western blot for HA-tagged mNeonGreen protein[45] (Fig. 3D). HEK293T cells were co-transfected with IRE1-XBP1 reporter along with either control or *H19*-overexpressing plasmid. At 16 hours post-transfection cells were treated with TG for 8 or 24 hours. Compared to control cells, *H19*-overexpressing cells exhibited a marked reduction in GFP fluorescence at both time points (Fig. 3E). Consistently, western blot analysis revealed decreased levels of HA-tagged mNeonGreen protein in *H19*-overexpressing cells (Fig. 3F). These results suggest that *H19* suppresses the activation of IRE1–XBP1 pathway during ER stress. To examine *H19*’s effect on the PERK-ATF4 signalling branch we used PERK activity reporter comprising *ATF4* 5′ UTR sequence, where ORF1 and ORF2 are used as translation initiation in non-stressful conditions. The phosphorylation of eIF2α by activated PERK stimulates translation initiation at ORF3 during conditions of ER stress. In PERK activity reporter, the fusion of HA-tagged-mScarlet-I and c-myc NLS coding sequence in frame with the first 84 nucleotides of ORF3 enables the detection of ATF4 translation by quantifying RFP fluorescence and/or western blot for HA-tagged-mScarlet-I protein[45] (Fig. 3G). HEK293T cells were co-transfected with PERK activity reporter along with either control or *H19*-overexpressing plasmid. Following treatment with TG for 8 or 24 hours, *H19*-overexpressing cells displayed increased RFP fluorescence intensity as compared to control cells (Fig. 3H). This observation was corroborated by western blot analysis, which showed elevated levels of HA-tagged ATF4 protein in *H19*-overexpressing cells (Fig. 3I). Together, these findings suggest that *H19* enhances activation of the PERK– ATF4 pathway during ER stress.

Next, we generated *H19*-overexpressing stable clones to investigate the impact of *H19* on UPR signalling. For this purpose, HEK293T cells were transduced with either lentivirus expressing *H19* and their corresponding control followed by puromycin selection. RT-qPCR analysis confirmed *H19* overexpression in 293T-H19 subclone, as indicated by a significant reduction in Ct values as compared to control (293T-Ctrl) subclone (Fig. 4A). Next, we assessed both protein and mRNA levels of downstream UPR targets in 293T-H19 and 293T-Ctrl subclones under ER stress conditions. Western blot analysis revealed that induction of XBP1s was attenuated in 293T-H19 cells as compared to 293T-Ctrl cells, indicating suppression of the IRE1-XBP1 axis of the UPR (Fig. 4B). During UPR conditions full length ATF6 (90 KDa) is transported to the Golgi apparatus to be processed by site 1 and site 2 proteases into cytosolic ATF6 (50 KDa) fragment, as such decrease in full length ATF6 protein can be used as a surrogate for its proteolytic processing. We observed that level of full-length ATF6p90 was reduced in both 293T-H19 and 293T-Ctrl cells at 4 hr of TG treatment, indicating proteolytic cleavage and activation of ATF6. However, the amount of full length ATF6 (90 KDa) was significantly reduced in 293T-H19 as compared to 293T-Ctrl cells suggesting prolonged activation of ATF6 (Fig. 4B). The basal and UPR-induced expression of phosphorylated eIF2α (P-eIF2α) and PERK were significantly higher 293T-H19 as compared to 293T-Ctrl cells (Fig. 4B). Next, we examined levels of UPR target genes (CHOP, HERP, GRP78, DNAJB9 and SEC24D) to determine if *H19* modulated their expression. RT-qPCR analysis revealed differential regulation of UPR target genes in response to TG-induced ER stress in 293T-H19 and 293T-Ctrl cells. The expression of all these genes (CHOP, HERP, GRP78, DNAJB9 and SEC24D) were upregulated in both 293T-H19 cells and 293T-Ctrl cells. The induction of ATF6 downstream genes (GRP78 and HERP) and PERK downstream gene (CHOP) was significantly higher in 293T-H19 as compared to 293T-Ctrl cells, whereas induction of XBP1 downstream gene (DNAJB9) was markedly attenuated in *H19*-overexpressing clones (Fig. 4C). Strikingly, both protein and transcript analyses consistently demonstrated that *H19* selectively upregulated the ATF6 and PERK pathways while suppressing the activation of IRE1-XBP1 pathway during ER stress. Importantly, these findings in *H19* stable clones mirrored the regulatory effects of *H19* observed in the reporter assays. Taken together, these results demonstrate that *H19* plays a specific role in modulating ER stress signalling, enhancing ATF6 and PERK activation while repressing IRE1 activity during ER stress.

**Figure 4:**
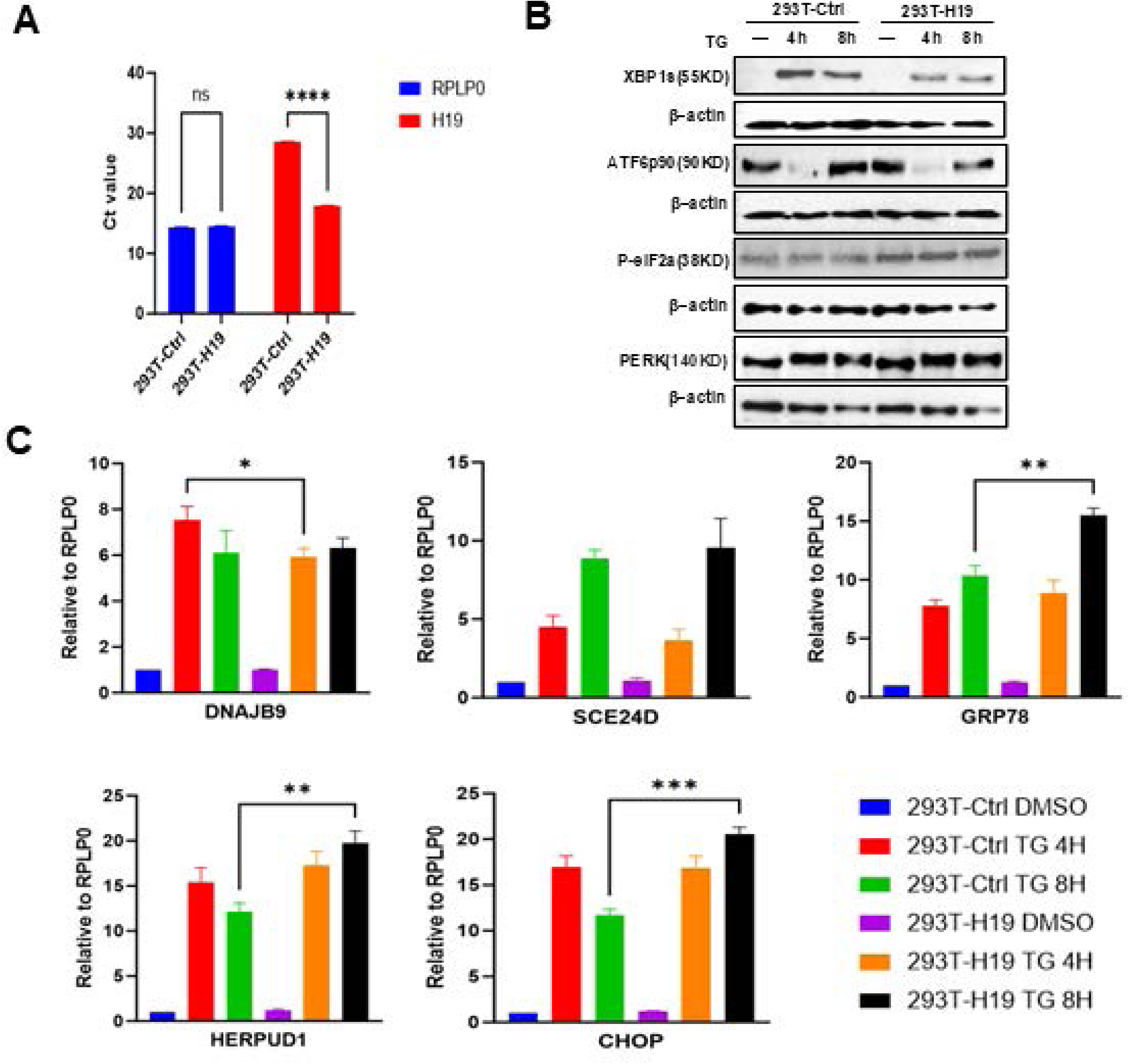
Effect of *H19* on expression of UPR-target genes. (A) RT-qPCR analysis of *H19* expression in HEK293T cells stably transduced with H19-overexpressing lentivirus (293T-H19) or control PLKO lentivirus (293T-Ctrl). (B) Stable 293T-H19 and 293T-Ctrl clones were treated with DMSO or thapsigargin (TG, 1 µM) for 4 or 8 hours. Total protein lysates were analysed by western blot to assess the expression of key UPR markers: XBP1s, ATF6p90, phosphorylated eIF2α (P-eIF2α), and PERK. β-Actin was used as a loading control. (C) Total RNA was extracted from the same cells and subjected to RT-qPCR to quantify the relative expression of UPR target genes: *DNAJB9*, *SEC24D*, *GRP78*, *HERPUD1*, and *CHOP*. Expression levels were normalized to *RPLP0*. Data represent the mean ± SD from three independent experiments. Statistical significance was assessed using an unpaired t-test:*P < 0.05; **P < 0.01; ***P < 0.001.

Next we determined the effect of *H19* on cell fate under ER stress conditions. For this293T-H19 and 293T-Ctrl cells were exposed to a wide range of BFA and TG and percentage of viable cells was measured at 48 h. These experiments revealed that 293T- H19 subclones exhibited significantly higher viability as compared to 293T-Ctrl under ER stress (Fig. 4A–B), suggesting a protective role for *H19*. We then directly assessed cell death via flow cytometry. Following 48-hour treatment with BFA and TG, *H19*- overexpressing clones showed significantly lower levels of cell death as compared to control clones (Fig. 4C–D), further supporting the notion that *H19* enhances cellular resistance to ER stress-induced apoptosis.

### *H19* expression is associated with poor outcome in TNBC

Breast is one of the few organs in which *H19* is not completely repressed after birth[46, 47]. *H19* is frequently upregulated in breast cancer tissues, and contributes to tumourigenesis through diverse molecular mechanisms, including epigenetic regulation, microRNA sponging, and modulation of signalling pathways[48–50]. *H19* has a multifaceted role in the progression of breast cancer, particularly in aggressive subtypes such as triple-negative breast cancer (TNBC). Univariate Cox regression analyses across various datasets (TCGA and METABRIC) demonstrated that elevated *H19* expression was associated with worse overall survival (OS), specifically in basal-like breast cancer (Fig. 6A). In the METABRIC dataset, high *H19* expression in basal-like patients was linked to poorer survival, with a hazard ratio (HR) of 1.97 (95% CI: 1.29– 3.00; p = 0.0017) (Fig. 6B). Similarly, TCGA data showed an HR of 3.26 (95% CI: 1.19–8.97; p = 0.022) for high *H19* expression in basal-like patients (Fig. 6C). These findings suggest that increased *H19* expression was associated with poor OS in basal-like breast cancer across multiple independent cohorts. Indeed, overexpression of *H19* in MDA-MB-231 cells has been reported to promote anchorage independent growth in soft agar and formation of larger tumours in SCID mice.

We reasoned that increased expression of *H19* may promote cancer progression by enabling cells to better adapt and survive the stressful conditions within the tumour microenvironment, which can activate UPR and trigger cell death. To evaluate the effect of *H19* on cell fate under conditions of ER stress, we generated stable *H19*- overexpressing subclones and the corresponding control in the TNBC cell line MDA-MB-231. RT-qPCR analysis confirmed a significant upregulation of *H19* expression in *H19*-overexpressing cells (MDA-H19) as compared to control (MDA-Ctrl) cells (Fig. 6D). We then evaluated stress-induced cell death in MDA-H19 and MDA-Ctrl cells. Flow cytometry analysis showed that MDA-H19 cells exhibited significantly reduced levels of cell death as compared to MDA-Ctrl cells (Fig. 6E-F). Together, these data suggest that *H19* overexpression confers resistance to ER stress-induced cell death most likely through modulating the UPR pathways.

## Discussion

*H19*, a well-characterized lncRNA, has been implicated in diverse biological processes, including development, cellular proliferation, and disease progression[11, 21, 32]. In cancer, *H19* exhibits context-dependent roles, functioning either as an oncogene or tumour suppressor, depending on the cellular environment and tumour type [11, 15, 51]. In this study, we show that *H19* expression was significantly decreased under ER stress conditions (Fig. 1). Our study further demonstrates that *H19* is downregulated by the UPR in a PERK-dependent manner (Fig. 2). *H19* participates in several important biological processes such as embryogenesis, tumourigenesis, osteogenesis, and metabolism, which are affected during the development of many diseases. The expression of *H19* is tightly regulated under normal conditions but becomes dysregulated in human diseases through combination of epigenetic, transcriptional, and signalling mechanisms[29, 52–54]. *H19* expression is dysregulated in diabetes by hypermethylation of *H19* locus[55]; cardiac hypertrophy and ischemic heart disease by inflammatory cytokines[56, 57]; hypoxia-induced upregulation in neurodegeneration and loss of imprinting and hypomethylation in cancers[54, 58] . Our results suggest a role of UPR in dysregulation of *H19* expression in human diseases where activation of UPR has been documented such as diabetes, cardiovascular diseases, neurodegeneration and cancer.

Cancer cells often face high levels of ER stress due to their rapid growth, metabolic reprogramming, and the harsh tumour microenvironment (including hypoxia, nutrient deprivation, and oxidative stress), which in turn activates UPR, a critical adaptive mechanism that helps cancer cells survive stressful conditions in tumour microenvironment [59–61]. While prolonged ER stress conditions can switch UPR to apoptotic signalling, cancer cells can evade UPR-induced cell death and hijack the UPR to support proliferation, metastasis, and chemoresistance[62]. For instance, in the early stages of tumour development, the UPR can suppress tumourigenesis by inducing apoptosis in cells experiencing excessive ER stress[63]. This protective mechanism ensures the elimination of damaged cells that could otherwise contribute to malignancy[64, 65]. However, in established tumours, the UPR often shifts to a pro-survival mode, enabling cancer cells to adapt to the harsh conditions of the tumour microenvironment. This adaptive response allows cancer cells to maintain their proliferative capacity and resist therapy-induced stress, making the UPR a key contributor to tumour progression and chemoresistance[66]. We found that H19 inhibits ER stress-induced cell death, suggesting that it may serve as a protective factor for breast cancer cells under ER stress conditions (Fig. 5, 6). Our results support the previous reports linking lncRNA to UPR regulation and cancer cell survival[67–69]. Indeed, *H19* expression has been reported play an important role in the induction of chemoresistance in several cancers. The *H19* expression impairs the efficacy of chemotherapy, endocrine therapy, and targeted therapy in breast cancer, liver cancer, lung cancer and colorectal cancer[30, 70, 71].

**Figure 5:**
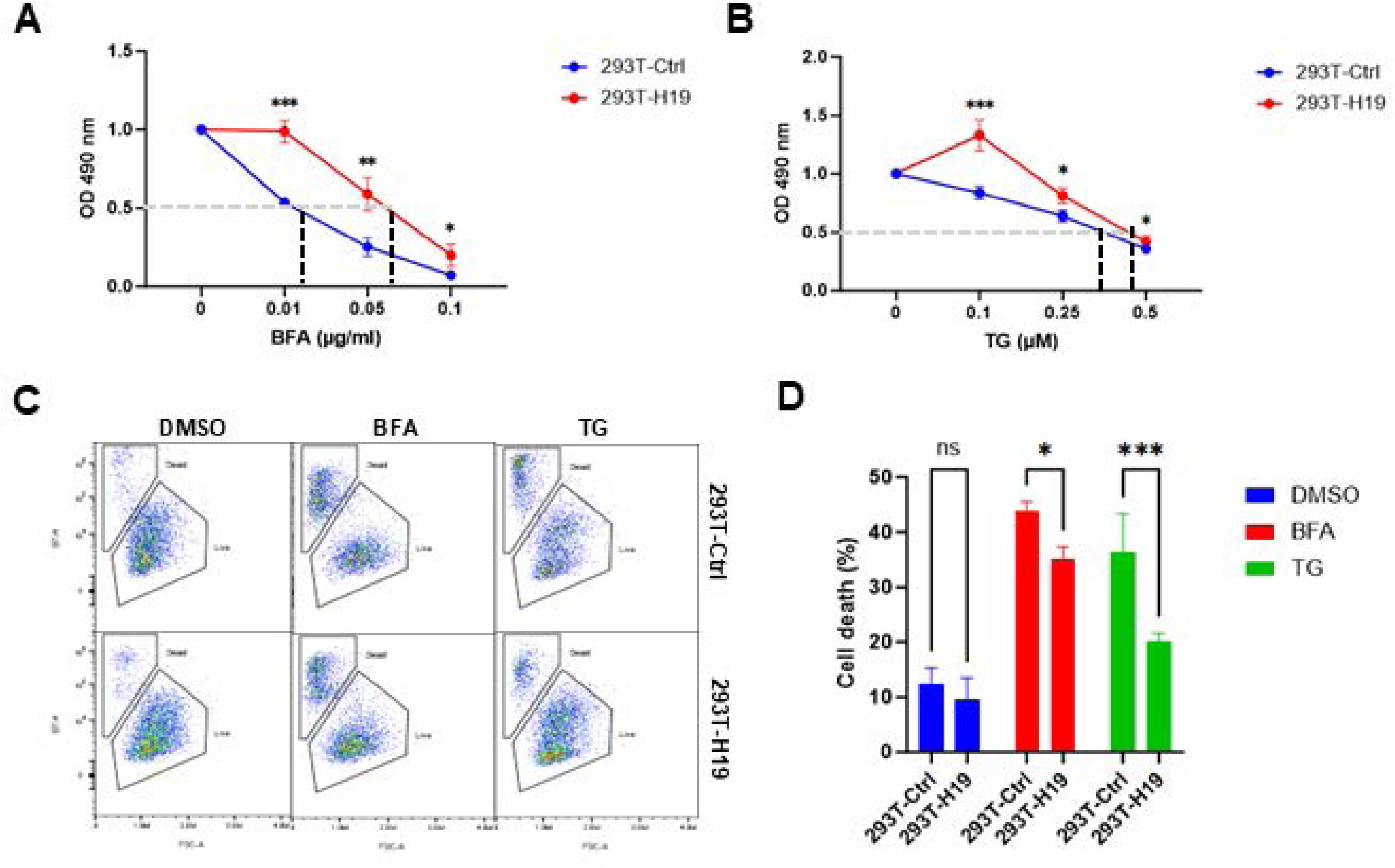
*H19* expression inhibits ER-stress induced cell death. (A–B) Cell viability of 293T-H19 and 293T-Ctrl subclones was evaluated by MTS assay following treatment with increasing concentrations of brefeldin A (BFA) (A) or thapsigargin (TG) (B) for 48 hours. Absorbance at 490 nm was measured, and fold change relative to untreated controls was calculated. Data represent mean ± SD from three independent experiments, each performed in four replicates. (C–D) Cells were treated with BFA (0.05 µg/ml) or TG (0.25 µM) for 48 hours, and cell death was assessed via propidium iodide (PI) staining and flow cytometry. (C) Representative flow cytometry plots and gating strategy. (D) Quantification of PI-positive (dead) cells. Unstained cells served as negative controls and PI-stained cells as positive controls. Data are presented as mean ± SD from three independent experiments. Statistical analysis was performed using an unpaired t-test:*P < 0.05; **P < 0.01; ***P < 0.001; ****P < 0.0001.

In our study, we also investigated the role of *H19* in regulating the UPR. Under conditions of ER stress, *H19* was found to enhance the signalling via the ATF6 and PERK arms while attenuate the signalling via the IRE1 arm of the UPR (Fig. 3, 4). The PERK arm of the UPR plays a dual role in determining cell fate: initially promoting survival by phosphorylation of eIF2α and reducing protein translation, leading to enhanced stress adaptation, but eventually triggering apoptosis under prolonged stress through CHOP activation. In our study, *H19* was found to potentiate PERK signalling, which suggests that it may reinforce the early cytoprotective phase of the response. The suppression of IRE1, which is often linked to pro-apoptotic signalling through JNK activation, could further prevent stress-induced cell death. Additionally, the activation of ATF6 by *H19* might facilitate DNA repair mechanisms and promote cell survival, as seen in colon cancer cells[72]. By modulating the UPR, *H19* attenuates ER stress-induced apoptosis in cancer cells, thereby promoting their survival. These findings indicate that *H19* plays a crucial role in shaping the ER stress response and UPR signalling to favour cancer cell adaptation under stressful conditions. *H19* could modulate UPR via several mechanisms – production of *miR-675*, acting as competitive endogenous RNA and miRNA sponge and interaction with chromatin modifiers and transcription factors. Further work is required to identify the key downstream effectors of *H19* that play a role in modulation of UPR signalling.

The *H19* is expressed during embryonic development and down-regulated after birth Expect in few tissues like mammary gland. There have been conflicting reports about its role in tumour initiation and progression, however accumulating data suggest that *H19* is one of the major genes in cancer. Several studies have shown that the *H19* is a prognostic biomarker in various human cancers. A meta-analysis based on 15 studies with 1584 patients and TCGA data for prognostic value of *H19* in various human cancers has shown that increased *H19* expression is associated with poor OS (HR= 1.62, 95% CI= 1.36-1.93, P<.001) in various cancers. The association of *H19* expression with poor OS was further validated by analysis using TCGA dataset consisting of 7462 cancer patients (HR= 1.12, 95% CI= 1.03-1.22, P < .05). Our results show that increased *H19* expression was unequivocally associated with poor OS in basal-like breast cancers from METABRIC and TCGA datasets (Fig. 6). Further, *H19* overexpression in MDA-MB-231 cells was reported to increase anchorage-independent growth and tumourigenicity *in vivo* but had no significant effect on the anchorage-dependent growth[48]. Our results show that *H19* overexpression MDA-MB-231 cells confers resistance to ER stress-induced cell death (Fig. 6).

**Figure 6:**
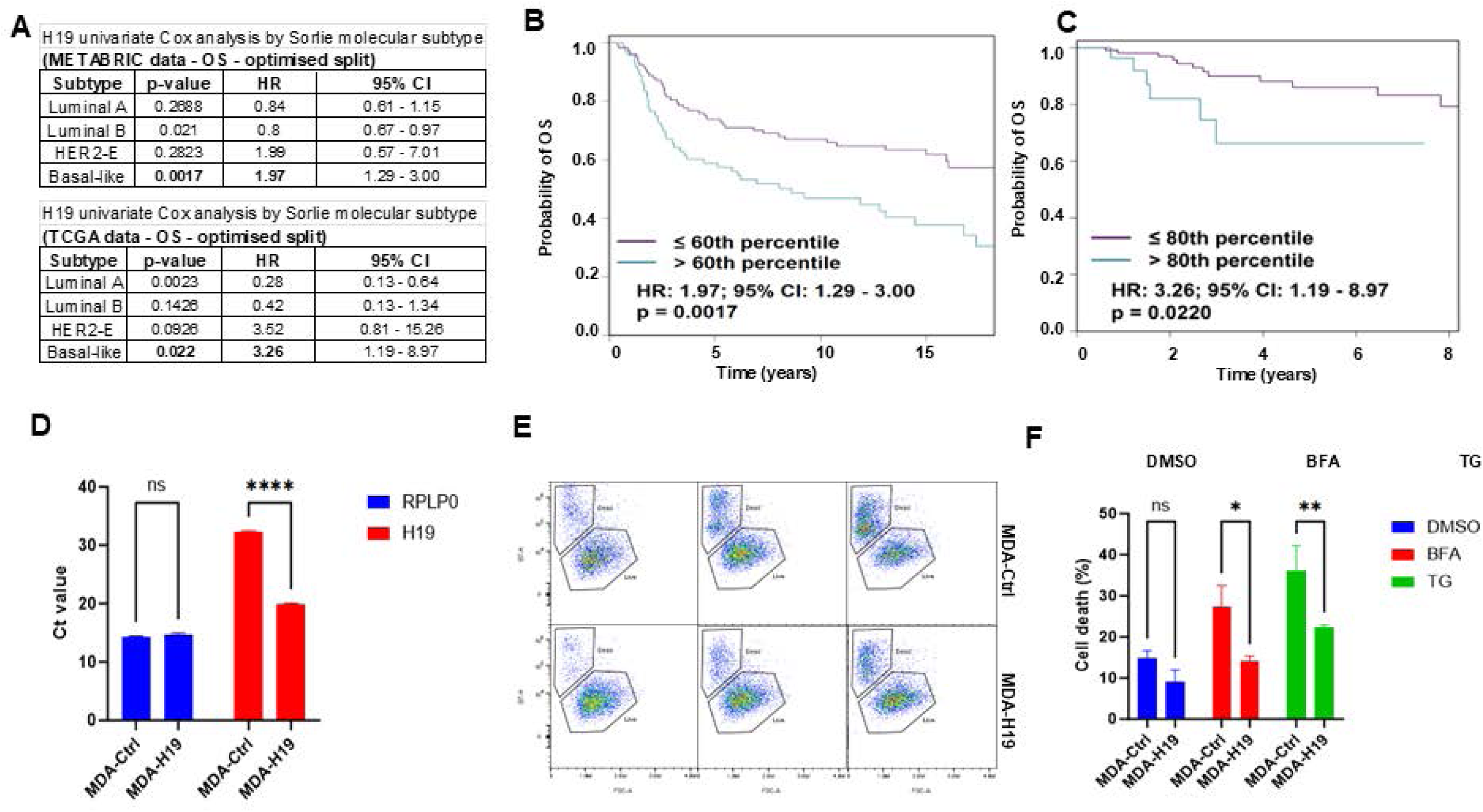
Association of *H19* expression with overall survival in triple-negative breast cancer. (A) Bioinformatic analysis of *H19* expression based on Breast Cancer Gene-Expression Miner v5.2, analysing all RNA-seq data according to RIMSPC subtypes. (B) The Univariate Cox analysis between *H19* expression levels and patient outcomes, Overall Survival (OS) across various breast cancer subtypes. (C-D) Kaplan-Meier survival estimates of *H19* expression in basal-like patients from METARIC data (C) and TCGA data (D). (E) *H19* expression was determined by RT-qPCR using total RNA from MDA-MB-231 *H19*-overexpressing (MDA-H19) and control (MDA-Ctrl) subclones. (F) MTS assays were performed for D1, D3, and D5 in MDA-H19 and control MDA-Ctrl subclones. Absorbance was measured at 490 nm, and fold change in absorbance relative to D0, 24 hours after cell plating, is shown. (C) MTS assays were performed after 48-hour treatment with increasing concentrations of BFA. Absorbance was measured at 490 nm, and fold change in absorbance relative to untreated is shown. Data presented as mean ±SD from three independent experiments. Each experiment was performed in four replicates. (H-I) Cells were subsequently treated with BFA (0.025 µg/ml) or TG (0.25 µM) for 48 hours. Gating strategy and representative flow plots are presented in (H), and statistical analysis of dead cell percentages is presented in (I). Data are presented as mean ± SD from three independent experiments. Statistical significance was determined using an unpaired t-test: *p < 0.05, **p < 0.01, ***p < 0.001, ****p < 0.0001.

Taken together our results reveal a pivotal role of *H19* in regulating cell fate during UPR. Considering our results, we speculate that the downregulation of *H19* may modulate cell growth and survival of TNBC by modulation UPR signalling in tumour microenvironment (Fig. 7). Given the ability of *H19* to modulate ER stress-induced apoptosis, targeting the UPR-H19 axis could represent a novel therapeutic strategy. Indeed, strategies to target lncRNAs that abrogate the expression of cancer-promoting lncRNAs such as *MALAT1* and *XIST* in breast cancer have been reported[73, 74]. Disrupting this interaction may sensitize cancer cells to ER stress-induced apoptosis, thereby limiting tumour growth and overcoming resistance to conventional therapies.

**Figure 7:**
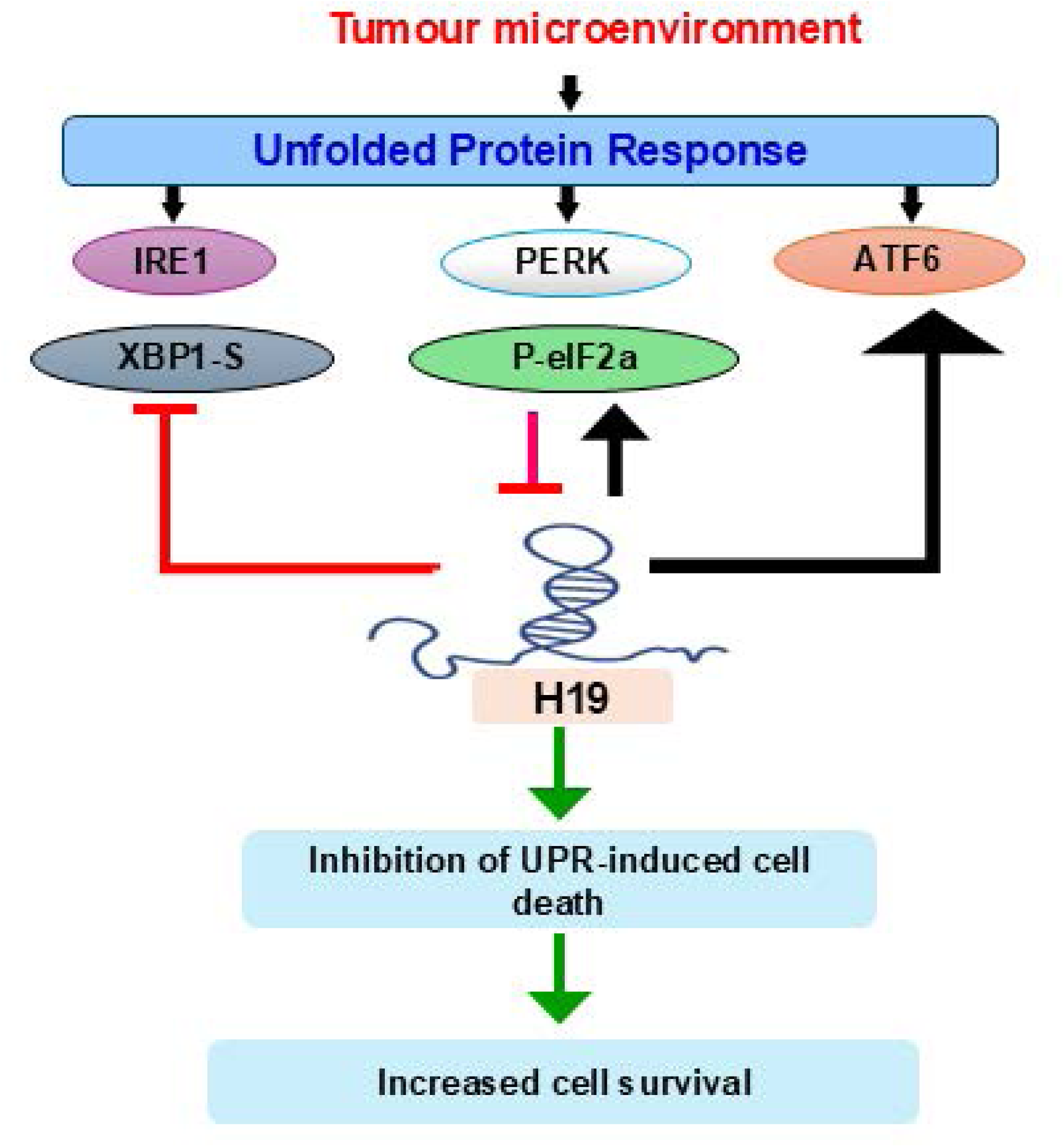
Graphical abstract. *H19* acts as a regulatory hub modulating the unfolded protein response (UPR) in breast cancer under endoplasmic reticulum (ER) stress. UPR activation leads to the downregulation of *H19* in a PERK-dependent manner. In a feedback loop, *H19* selectively upregulates the ATF6 and PERK branches while suppressing the IRE1/XBP1 axis of the UPR. This reprogramming of UPR signalling by *H19* enhances cell survival and attenuates ER stress-induced apoptosis, particularly in triple-negative breast cancer (TNBC). *H19* overexpression confers resistance to ER stress-inducing agents (e.g., thapsigargin and BFA), while its depletion sensitizes cells to UPR-driven and therapeutic stress. These findings underscore *H19* as a potential modulator of UPR plasticity and a promising therapeutic target for sensitizing breast cancer cells to ER stress-mediated cell death.

## Materials and Methods

### Cell Culture and Drug Treatment

Breast cancer cell lines representing different subtypes—MCF7, T47D, BT474, and MDA-MB-231—were obtained from ECACC (Salisbury, Sussex, UK). HEK293T cells were sourced from the Indiana University National Gene Vector Biorepository (Indianapolis, IN, USA). MCF7, BT474, MDA-MB-231, and HEK293T cells were cultured in DMEM supplemented with 10% foetal calf serum, 100 U/ml penicillin, and 100 μg/ml streptomycin at 37°C with 5% CO₂. T47D cells were maintained under the same conditions but in RPMI-1640 medium. To induce ER stress, cells were treated with brefeldin A (BFA; Tocris, Cat #1231) or thapsigargin (TG; Tocris, Cat #1138) at the specified concentrations and time points. To inhibit specific UPR arms, cells were treated with Ceapin A7 (ATF6 inhibitor; Bio-Tech, Cat #6955), STF083010 (IRE1α inhibitor; Merck Millipore, Cat #SML0409), or GSK2606414 (PERK inhibitor; Merck Millipore, Cat #516535). PERK activation was selectively induced using the EIF2AK3 activator CCT020312 (Merck Millipore, Cat #324879).

### RNA Extraction, cDNA Synthesis, and RT-qPCR

Total RNA was extracted using TRIzol™ Reagent (Thermo Fisher Scientific, Cat. #15596026) according to the manufacturer’s protocol. Real-time quantitative PCR (RT-qPCR) was performed using a two-step process. First, cDNA was synthesized from 1– 5 µg of total RNA using the HiScript® II 1st Strand cDNA Synthesis Kit (Generon Reagents Distributor Limited, Cat. #R211-01). Gene expression was then analyzed using the resulting cDNA and predesigned PrimeTime® qPCR assays (Integrated DNA Technologies, Belgium). For miRNA-qPCR, specialized primers—such as stem-loop primers or modified universal primers—were employed to ensure specific and efficient amplification, using the TaqMan™ MicroRNA Kit (Thermo Fisher Scientific, Cat. #4427975). Relative expression levels were calculated using the 2^−ΔΔCT method.

### Protein Extraction and Western Blot Analysis

Cells were washed once with ice-cold PBS and lysed in RIPA buffer following the indicated treatment times. Protein concentration was determined using the Bradford assay. Equal amounts of protein were mixed with Laemmli’s SDS-PAGE sample buffer, boiled at 95°C for 5 minutes, and loaded onto an SDS-polyacrylamide gel for electrophoresis. Proteins were then transferred onto a nitrocellulose membrane and blocked with 5% milk in PBS containing 0.05% Tween-20 (PBST). The membrane was incubated overnight at 4°C with primary antibodies against ATF6 (Abcam, Cat #ab122897), spliced XBP1 (BioLegend, Cat #619502), PERK (Cell Signaling, Cat #C33E10), phospho-eIF2α (Cell Signaling, Cat #9721S), β-Actin (Sigma, Cat #A- 5060), HA-Tag (Cell Signaling, Cat #MA1-12429), and/or Tri-methyl-histone H3 (K4) (Cell Signaling, Cat #9727S). After three washes with PBST, the membrane was incubated with the appropriate horseradish peroxidase-conjugated secondary antibody for 2 hours at room temperature. Signals were detected using the Pico PLUS Chemiluminescent Substrate (BioSciences Limited, Cat #34580).

### Transient Transfection

Human *H19* (ENST00000412788) full-length cDNA was cloned into the control vector pLenti4/V5 (Life Technologies), generously provided by Reinier from the Institute of Cardiovascular Regeneration, Goethe University, Frankfurt, Germany. Empty pLenti4/V5 vectors were used as controls. The HA-tagged ATF4 (Addgene, Cat #115969) and XBP1 (Addgene, Cat #115968) reporter plasmids were purchased from Addgene. For transfection in HEK293T cells, 1 µg of plasmid DNA was mixed separately with 6 µL of JetPEI (VWR, Cat #101-10N) in 100 µL of 150 mM NaCl. For transfection in MCF7 cells, 1 µg of plasmid DNA was mixed separately with 2 µL of TurboFect (Fisher Scientific, Cat #15325016) in 100 µL of serum-free Opti-MEM (Biosciences, Cat #11058021). The transfection reagent and plasmid DNA solutions were incubated separately at room temperature for 10 minutes, then combined and incubated for an additional 20 minutes. The resulting 200 µL DNA complex was then added to 2 mL of fresh complete DMEM for 293T cells or Opti-MEM for MCF7 cells. After 4 hours, Opti-MEM was replaced with complete DMEM.

### Generation of Stable Cell Lines

*H19*-overexpressing clones were generated via lentiviral transduction. Lentiviral stocks were produced in HEK293T cells using PMD2.G and psPAX as packaging plasmids, along with the *H19* overexpression vector (Addgene, Cat #200835). Empty PLKO vectors were used as controls. MDA-MB-231 Cells were transduced with *H19*- overexpressing lentiviral or PLKO lentiviral for 24 hours, after which positive clones were selected using puromycin-containing media (1 µg/mL) for one week.

### Luciferase Reporter Assay

The 5×ATF6-pGL3 (ATF6 pathway reporter) contains five copies of ATF6-binding sites (CTCGAGACAGGTGCTGACGTGGCGATTC) cloned into pOFlucGL3, upstream of the c-fos minimal promoter (−53 to +45 of the human c-fos promoter). This reporter plasmid was obtained from Addgene (Cat #11976). 24 hours post-transfection, cells were treated with TG for 24 hours. Firefly luciferase and Renilla luciferase activities were measured 48 hours after transfection using the Lucetta™ Luminometer (Lonza). Firefly luciferase activity was then normalized to Renilla luciferase activity for data analysis.

### Flow Cytometry Analysis

Cells were plated in 6-well plates. After 24 hours, cells were treated with either vehicle or the required compounds for the indicated time points. The media was collected into a separate 15 mL tube, and cells were harvested by trypsinization. Following centrifugation at 200 g for 5 minutes at 4°C, the media was discarded, and cells were washed once with ice-cold PBS. The cells were then resuspended in fluorescent-activated cell sorting buffer. Gating was performed using propidium iodide (PI) unstained cells and positive control cells stained with PI (0.25 µg/mL). The percentage of live or dead cells was determined using the Cytek Northern Lights 2000 Flow Cytometer, and data analysis was performed using FlowJo v10.9.0.

### MTS assay

1000 cells are seeded in a 96-well plate at an optimized density in 150 µL of complete culture medium. After 24 hours of incubation, cells may be treated with compounds for a specific duration or at multiple time points (D0, D1, D3, D5) to track cell viability or proliferation. At each testing time point, 20 µL of MTS solution (2 mg/mL 3-(4,5- dimethylthiazol-2-yl)-5-(3-carboxymethoxyphenyl)-2-(4-sulfophenyl)-2H-tetrazolium (MTS) (Promega, Cat #G3582) and 0.9 mg/mL phenazine methosulfate (PMS), mixed at a 10:1 ratio) is added to 100 µL of medium in each well. The plate is then incubated at 37°C for 1.5 hours. Control wells containing only medium and MTS reagent (without cells) are used for background subtraction.

### Statistical Analysis

Data were analyzed using GraphPad Prism 10.1. Results are presented as mean ± SD from three independent experiments, unless otherwise stated. The p-value was determined using an unpaired t-test between independent groups, and results with p < 0.05 were considered statistically significant.

## Author contributions

S.G: Conceptualization, Supervision, Methodology, Visualization, Formal Analysis, Writing – Original Draft.

W.L: Investigation, Visualization, Data Curation, Formal Analysis, Writing – Original Draft.

A.G: Conceptualization, Visualization, Supervision, Formal Analysis, Writing – Review and Editing.

M.K: Conceptualization, Resources, Writing – Review and Editing, Funding Acquisition.

All authors have read and approved the final manuscript.

## Acknowledgements

W.L. was supported by the China Scholarship Council (No. 202106370068). S.G received research funding from National Breast Cancer Research Institute under Galway University Foundation grant number FY24001. We would like to express sincere gratitude to members of S.G laboratory Afrin Sultana, Wenyuan Zhao, Qian Xu, and Dong Ning for valuable discussions throughout this project.

## Abbreviations

ATF4: activating transcription factor 4
ATF6: activating transcription factor-6
Bcl-2: B-cell lymphoma 2
BFA: Brefeldin A
BIM: Bcl-2-interacting mediator of cell death
ceRNAs: competing endogenous
RNAs DR5: death receptor 5
EMT: epithelial–mesenchymal transition
ER: endoplasmic reticulum
ERAD: ER-associated degradation
GOLGA2P10: golgin A2 pseudogene 10
GRP78: glucose-regulated protein 78
IRE1α: inositol requiring enzyme1α
lncRNAs: Long non-coding RNAs
OS: overall survival
PERK: protein kinase RNA-like endoplasmic reticulum kinase
PI3K/AKT: phosphoinositide 3-kinase/protein kinase B
TG: Thapsigargin
TNBC: triple-negative breast cancer
UPR: unfolded protein response
XBP1: X-box binding protein-1
XBP1s: spliced X-box binding protein-1

---

**Supplementary Figure 1.**
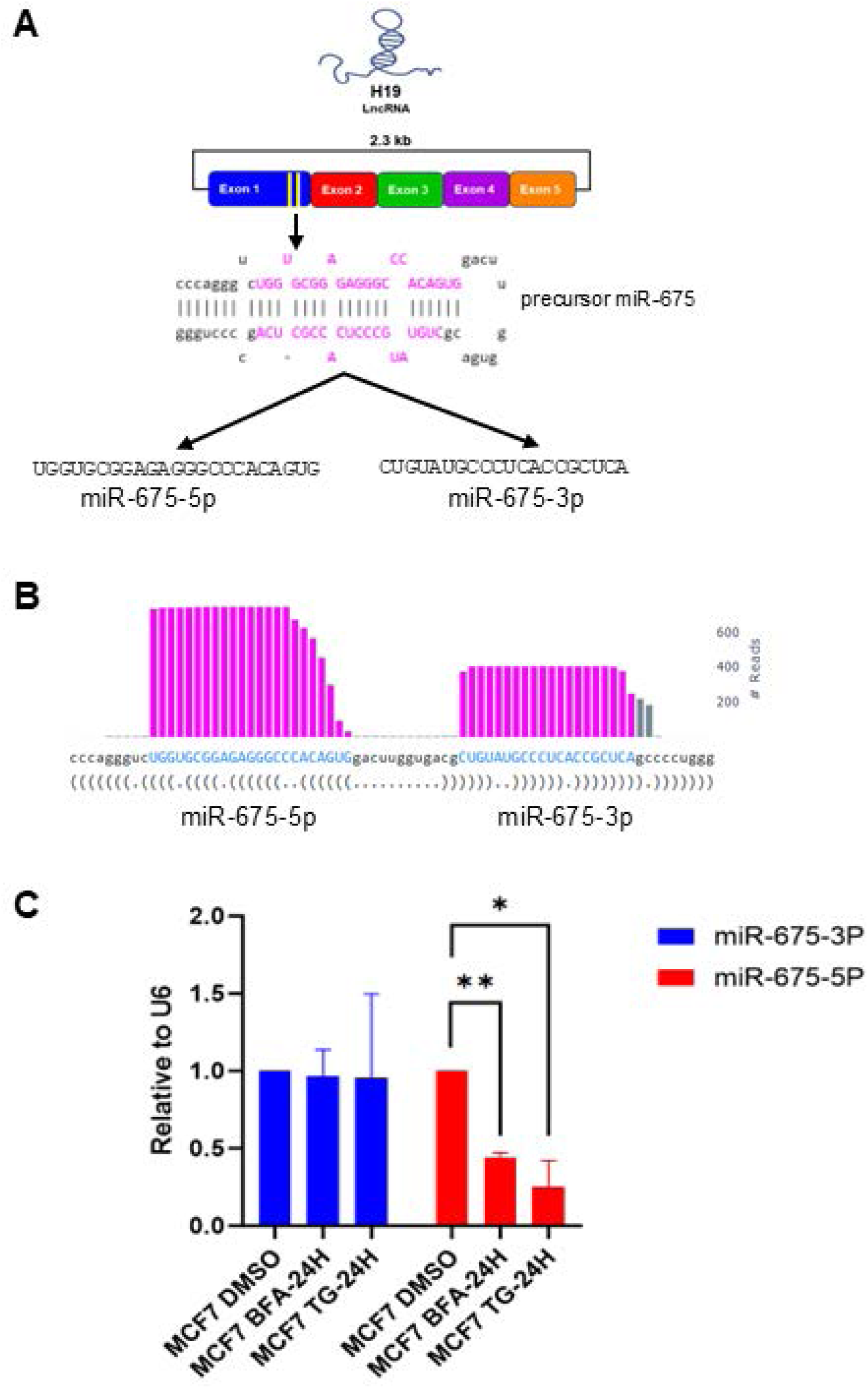
Expression of miR-675-5p is downregulated by UPR. **(A)** Schematic representation of biogenesis and sequence of miR-675-5p and miR-675-3p derived from exon-1 of the H19 gene. (B) The relative number of reads for miR-675-5p and miR-675-3p from miRbase are shown. (C) Expression levels of miR-675-3p and miR-675-5p in MCF7 cells following 24-hour treatment with either DMSO, BFA (0.5 µg/ml), or TG (1 µM). Expression levels were quantified relative to U6 (a normalization control). Statistical significance was determined using an unpaired t-test compared with DMSO-treated cells. *P < 0.05, **P < 0.01.

